# An Assistive Computer Vision Tool to Automatically Detect Changes in Fish Behavior In Response to Ambient Odor

**DOI:** 10.1101/2020.09.01.277657

**Authors:** Sreya Banerjee, Lauren Alvey, Paula Brown, Sophie Yue, Lei Li, Walter J. Scheirer

## Abstract

The analysis of fish behavior in response to odor stimulation is a crucial component of the general study of cross-modal sensory integration in vertebrates. In zebrafish, the centrifugal pathway runs between the olfactory bulb and the neural retina, originating at the terminalis neuron in the olfactory bulb. Any changes in the ambient odor of a fish’s environment warrants a change in visual sensitivity and can trigger mating-like behavior in males due to increased GnRH signaling in the terminalis neuron. Behavioral experiments to study this phenomenon are commonly conducted in a controlled environment where a video of the fish is recorded over time before and after the application of chemicals to the water. Given the subtleties of behavioral change, trained biologists are currently required to annotate such videos as part of a study. This process of manually analyzing the videos is time-consuming, requires multiple experts to avoid human error/bias and cannot be easily crowdsourced on the Internet. Machine learning algorithms from computer vision, on the other hand, have proven to be effective for video annotation tasks because they are fast, accurate, and, if designed properly, can be less biased than humans. In this work, we propose to automate the entire process of analyzing videos of behavior changes in zebrafish by using tools from computer vision, relying on minimal expert supervision. The overall objective of this work is to create a generalized tool to predict animal behaviors from videos using state-of-the-art deep learning models, with the dual goal of advancing understanding in biology and engineering a more robust and powerful artificial information processing system for biologists.

## Introduction

For many behavioral experiments in neuroscience, it is now a necessity to analyze large collections of digital video data captured in controlled or semi-controlled circumstances, where some degree of variation in the recorded scenes is expected. Examples include experiments for different animal model systems like fish, rodents, primates, drosophila, and worms [3, 8, 23, 25, 35, 39, 44], which study how these animals work together in groups or alone in response to certain stimuli. The stimulus can be a naturally occurring aspect of the environment (e.g., natural light), but is more often artificially induced in a laboratory setting. For example, the addition of odorants into water to understand the interplay between the olfactory and visual systems of a fish.

When looking for a change in behavior, the first step in analyzing video data is tracking the movement of individual animals across a sequence of recorded frames. This can be done algorithmically through the use of target detection and tracking methods from computer vision. Most of the tracking methods that are commonly deployed for this purpose are fully automatic [2, 10, 41, 42, 49], although there are some exceptions [31] where tracking requires some form of human intervention. The second step of the process is classifying a change in animal behavior. For this, automatic approaches have been suggested in the literature [10, 19, 23, 26, 33, 36, 40, 46], but there is no generally applicable approach as there is with target detection and tracking, and many scientists opt to perform this stage by hand. Thus even with partial automation, the video annotation process still requires extensive labor and time. Here we propose a new assistive vision tool that combines a novel generalized tracking algorithm based on deep learning with a behavior-specific classifier to better automate the video annotation process for behavioral experiments that look for distinct changes in animal behavior.

To develop and validate the proposed video analysis tool, we look at a specific case study of how olfactory signals affect vision through the centrifugal pathway in fish. For this study, we make use of videos of experimentation that behaviorally demonstrate this interplay between the visual and olfactory systems. The zebrafish (*Danio rerio*) is a common animal model system for studying multi-modal sensory integration. This is because it shares a high evolutionary proximity to mammals [12]. Zebrafish possess a prominent centrifugal pathway, running between the olfactory bulb and the neural retina, referred to as the olfacto-retinal centrifugal pathway (the ORC pathway) [24, 30]. The ORC pathway originates from the terminalis neurons (TNs), embedded in the olfactory bulb. Each olfactory bulb contains approximately 20-30 TNs. The TNs project axons to large brain areas, and through the optic nerve, some of the TN axons enter the retina where they extend in the interplexiform layer along the board of the inner nuclear layer. While propagating in the retina, the TN axons are branched, projecting to the inner nuclear layer and synapsing with dopaminergic amacrine cells.

The increased visual sensitivity in zebrafish due to odorants has been an active area of study for many years. Insights from relatively recent research [24, 30] have shown that the function of the ORC pathway is regulated by the olfactory input. Huang et al. [24] demonstrated how the visual sensitivity in zebrafish is increased in the presence of olfactory signals whereas disrupting the ORC pathway impairs visual function. They considered single-unit retinal ganglion cells for those experiments. Banerjee et al. [4] modeled this phenomenon by assuming the importance of extreme cellular responses in the modulation of sensory systems and applying the statistical extreme value theory.

Our experiments use a behavioral assay based on the visually-mediated escape responses of zebrafish. The experimental setup consists of a rotating cylinder and a stimulus, in the form of white paper marked with a black segment attached to the surface of the rotating cylinder. The fish is allowed to swim freely in the cylinder before and after the application of an odorant. It reacts to the stimulus when visible. The entire setup is illuminated by a light source from above, whose light intensity can be adjusted by changing neutral density filters. Additionally, the fish’s visual threshold is recorded using light filters to find the minimum amount of light required to produce an observable, visually mediated escape response. This swimming behavior is captured as a video via an infrared camera, with biologists subsequently performing data analysis from the video recordings by hand.

Typically, a fish swims slowly along the wall of the container in either clockwise or counterclockwise direction. However, when challenged by the black segment rotating outside the container, the fish displays a robust escape response, i.e., it turns and swims in the opposite direction of the stimulus. This sudden change in behavior can be attributed to the addition of an odorant in the water. As a result of its appearance, the fish becomes more aware of its surroundings due to the ORC pathway operating more effectively at a lower light intensity. Before the development of the proposed tool, collected videos had to be manually assessed by trained biologists to determine whether or not this behavior change was present.

Recent advances in computer vision technologies have allowed us to build a tool that immediately benefits this type of scientific experimentation. Our tool extracts meaningful information such as a fish’s trajectory using state-of-the-art deep learning and supervised classification methods to automate the prediction of behavioral changes when the sensory integration system is instantiated. It requires no human supervision in operation. Additionally, it can be generalized to a number of experiments that involve studying motion-based animal behavior. To date, this is the only tool that is able to automatically predict changes in behavior in zebrafish due to odor stimulation. In order to study the utility of this tool for behavioral experiments, we compare its output with human judgements from an attempt at crowdsourcing the video annotation. Based on the user study, the tool has been found to be accurate by a large margin of 50%, compared to humans.

To summarize, the contributions of this article are:

- A new fully automatic end-to-end tool for analyzing behavioral changes in fish that drastically increases data analysis throughput, and minimizes errors due to human bias and inattentiveness. The software is available as an open source package on GitHub.
- A detailed evaluation of the effectiveness and usability of the tool based on verified ground-truth annotations from experts, as well as a comparison to human annotations produced via crowdsourcing.

## Results

The proposed tool is a computational pipeline consisting of five stages, each of which is discussed in detail in the Methods section. Raw video showing the movement of a fish in the water after the application of chemicals is used as input to the tool. The tool outputs a binary “yes” or “no” decision indicating whether or not the fish exhibited behavioral change. Since the behavioral changes in fish are primarily studied by biologists through their swimming patterns, we use automatic target detection and tracking side-by-side to generate a specialized trajectory image from the raw video. Fig. 1 shows the process used to generate a trajectory image from experimental data. Compressed trajectory images are then used as input to a behavior classifier. Fig. 2 shows how the classification process follows the generation of trajectory images. In this section we describe two evaluations to validate the trajectory image quality and classification output.

**Figure 1.**
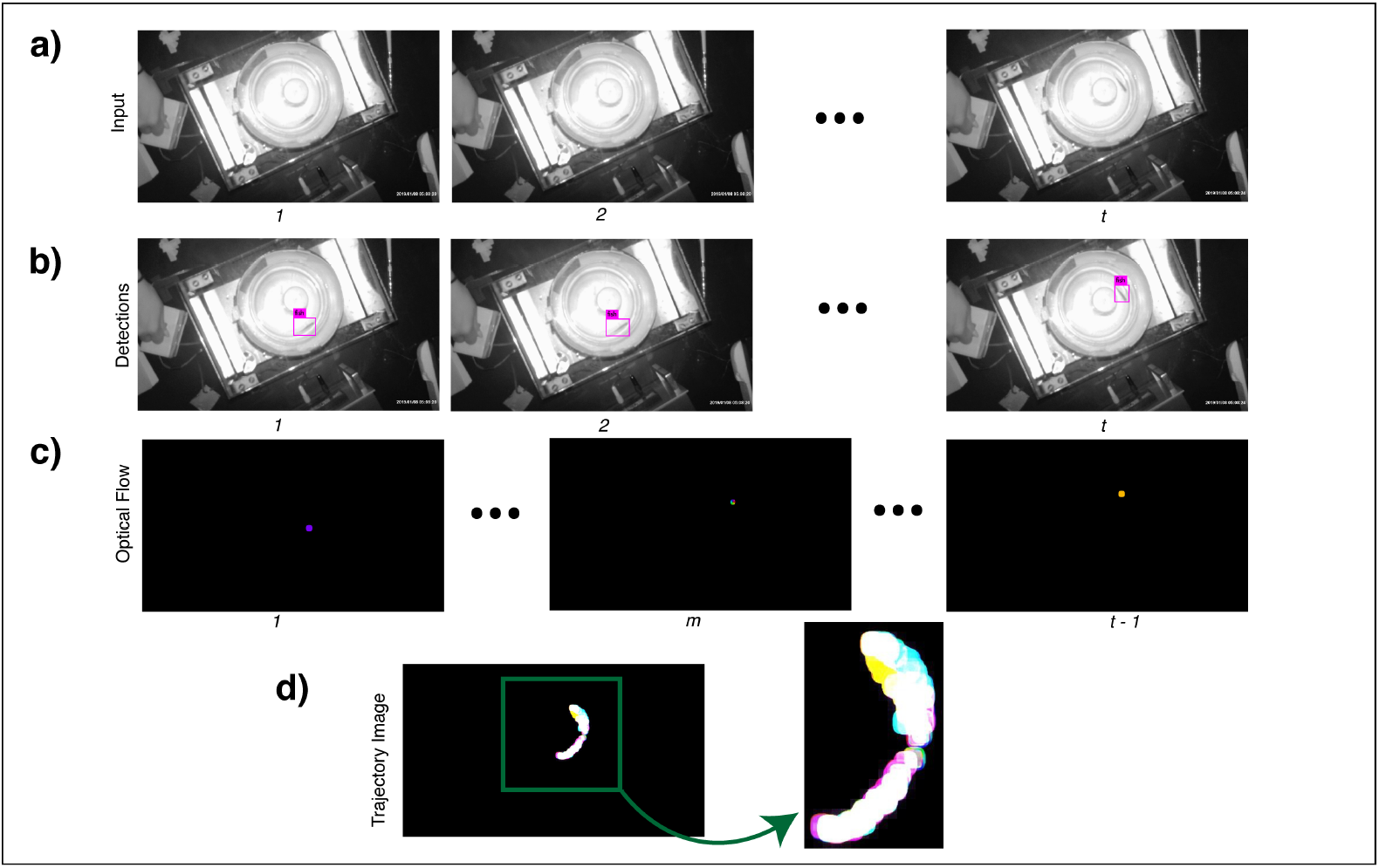
Process used to generate a trajectory image from experimental data. This figure illustrates the first two steps, fish detection and tracking, of the software pipeline that forms the proposed tool. The process includes (a) Selection of raw input video frames; (b) Automatic detection of a fish within each video frame; (c) Tracking of the fish via optical flow; (d) Creation of a trajectory image combining the optical flow output of the video frames, which is provided to an autoencoder for compression. We use the latent representation from the autoencoder for classification. That process is shown in Fig. 2. The numbers 1, 2,…, *t* just beneath the images stand for different timestamps of the video frames. Since the optical flow algorithm operates on two consecutive frames, the total number of frames after processing by the algorithm is *t* − 1.*Created with Adobe Illustrator CC Version 22.1.*

**Figure 2.**
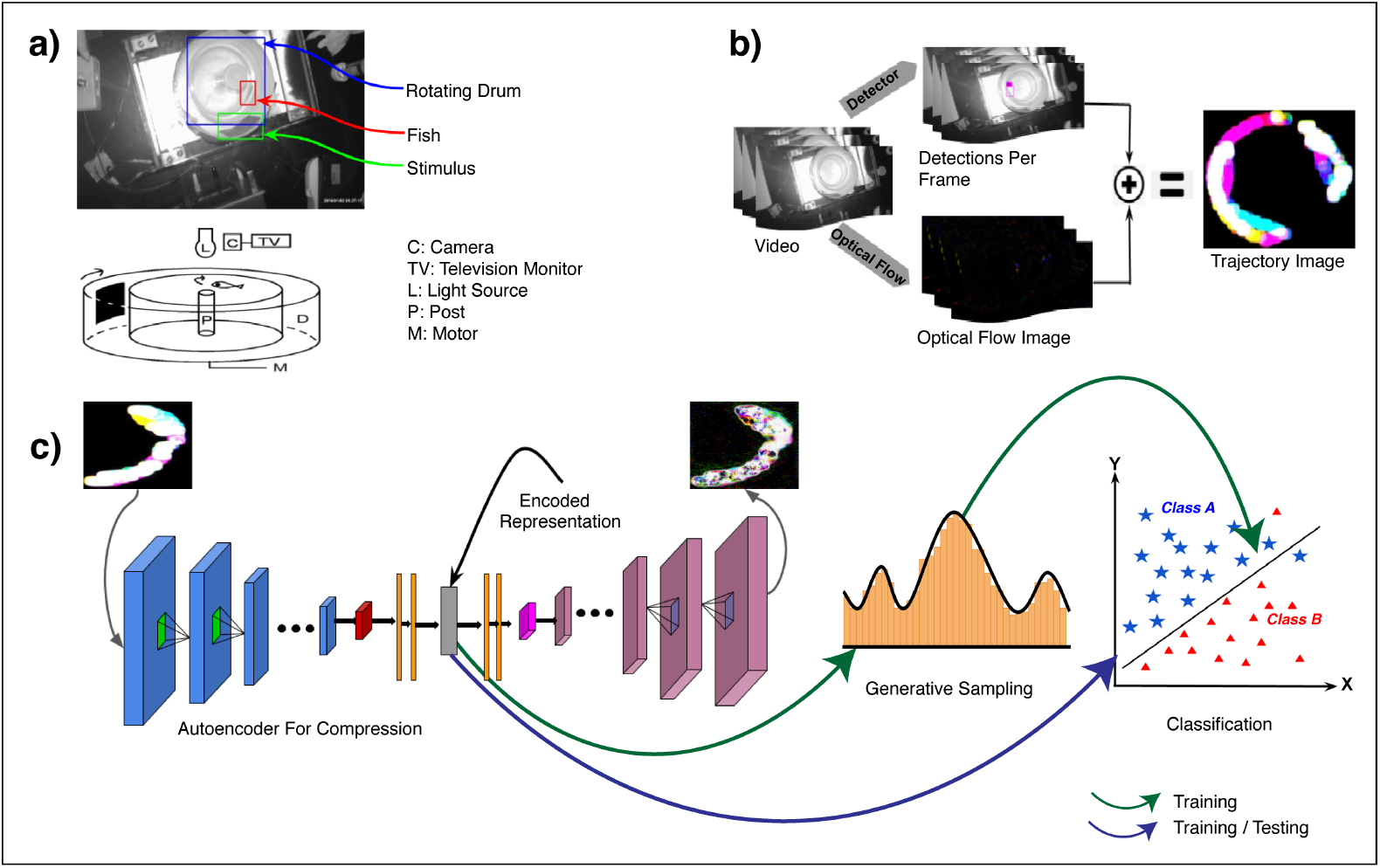
An overview of the behavioral experiments and tool to analyze them. **(a)** Experimental setup for recording behavioral visual sensitivity in zebrafish in response to olfactory and TN stimulation. The drum rotation in the lower diagram is clockwise, and the direction of the swimming fish is initially counterclockwise. The fish displayed escape responses to the approach of the black segment. Upon the black segment coming into view, a fish will immediately turn and swim away (in the clockwise direction in this example). Abbreviations used in the lower diagram: C, camera; D, rotating drum; L, light source; M, motor; P, post; TV, television monitor. **(b)** The process for generating trajectory images for zebrafish from videos. This shows how the first two steps of the overall pipeline (see Fig. 1) are combined to form a trajectory image. We use automatically detected regions of interest to create a mask for the fish such that only the pixels representing the fish in the tank are illuminated for dense optical flow estimation. All the frames are combined thereafter to generate a single trajectory image for the entire video. **(c)** Data compression using autoencoders, generative sampling and a binary classifier for behavior analysis for fish. This shows how the remaining three steps of the overall pipeline fit together. Since the raw features from the trajectory images can be high-dimensional, we use compression via autoencoders to limit the dimensionality of trajectory images. The encoded representations can be used as-is for classifier training and testing, or as priors for generative sampling before training a classifier.*Created with Adobe Illustrator CC Version 22.1.*

With respect to the dataset used for the evaluations, we collected 46 escape response videos from wet-bench experiments, as well as a single video representing zebrafish mating behavior. Out of these we reserved four escape response videos that were annotated by trained biologists and 300 frames from the mating video to create the fish detection dataset. Combining the video frames from the escape videos with the frames from the mating video, 1226 frames were available for training a zebrafish detector. Since we are only interested in detecting a single class (the presence of a zebrafish), this number of frames was sufficient. Out of the remaining 42 wet-bench videos, we utilized 31 videos to create a total of 160 short videos for training and testing behavior classifiers by splitting each video into clips that were four seconds in length. The time duration for each clip was suggested by biologists as being enough to notice changes in behavior in fish. Each four second clip was annotated by biologists to note the time of the onset of behavioral changes after the application of odorants in the water. We reserved 10 of the remaining videos for the human comparison study.

### Trajectory Image Evaluation

To evaluate the target (i.e., fish) detection performance in a quantitative manner, Mean Average Precision (mAP) at Intersection over Union (IoU) in the intervals [0.5, 0.75, 0.9] is used. The mAP evaluation follows the same protocol from the PASCAL VOC object detection evaluation regime [13], which is a standard in computer vision, except for a single modification introduced in IoU. IoU is the part of the evaluation that assesses the quality of the bounding boxes drawn around a detected target. Unlike PASCAL VOC, we evaluate mAP at different IoU intervals to account for different light intensity and experimental conditions. We consider predicted output to be a “true match” when it shares the same label as the ground-truth and has an IoU ≥ 0.5, 0.75, 0.90. The average precision (AP) for each class is calculated as the area under the precision-recall curve. Since we are only interested in detecting fish, i.e., a single class, mAP is equivalent to AP. We achieve a mAP score of 72.17% at IoU 0.5, meaning 72.17% of the time the fish was correctly detected with each predicted bounding box overlapping with the ground-truth pixels of the target by at least 50%. An example of a successful detection is shown in Supp. Fig. 1(A). For the comparatively difficult IoU intervals of 0.75 and 0.90, the mAP scores were relatively low, 34.12% and 0.40% respectively, due to near dark or low light intensity and false positive detections caused by reflections. Supp. Fig. 1(B) shows one such detection failure case due to a reflection on the surface of the tank. In that figure, one can see that there is a single fish swimming, which is correctly shown by the bounding box on the left. The detector fails though, as it identifies the reflection of the fish as a separate entity, making the count of fish two instead of one. Mis-detections or false positives do not necessarily pose a problem for the tool, as such mistakes tend to occur in unrelated frames amongst hundreds of frames with correct detections in a single video. We use these mistakes to our advantage when training the classifier portion of the tool, as they provide examples of noise that must be tolerated for generalization purposes.

The trajectory images generated as a result of detection and optical flow are very large feature-wise, in comparison to the number of videos we have from the biological experiments. As a result, we compress the images to reduce dimensionality via autoencoders (see the Methods section below for details of this process). Supp. Fig. 2 shows example reconstructed outputs from different autoencoder models. Higher dimensionality always warrants better reconstruction. However, to balance feature dimensionality with the number of available videos, we use 64 dimensions.

### Behavioral Classification Evaluation

To quantitatively evaluate the behavior classification performance of the proposed tool against human judgement, we use four metrics: Accuracy, Precision, Recall, and F1 Score. The data for this experiment came from the wet-bench experiment videos showing the escape response of zebrafish due to the addition of odorants into the water. Accuracy simply measures the ratio of correctly predicted observations to the total observations. Note that a higher accuracy does not always signify a better model. Thus we also use precision, which is the ratio of correctly predicted positive observations to the total predicted positive observations, and recall, which is the ratio of correctly predicted positive observations to all observations in the actual class. F1 score measures the weighted average of the precision and recall scores. The results are summarized in Table 1. As can be seen from the table, none of the classifiers achieve an accuracy greater than 65%. The non-expert human accuracy, on the other hand, is very low (34%), meaning that the task is very hard and cannot be outsourced to non-experts. The low accuracy score for classifier performance can be attributed to the fact that we were operating with a small number of videos, just 160, that could be used as training data for the system. While this is a sufficient number of videos for the manual data analysis necessary to study the effect at hand, it is, by current machine learning standards, rather limited. To address this, we turn to a data augmentation strategy that uses the available videos as a basis for generative sampling.

**Table 1.**
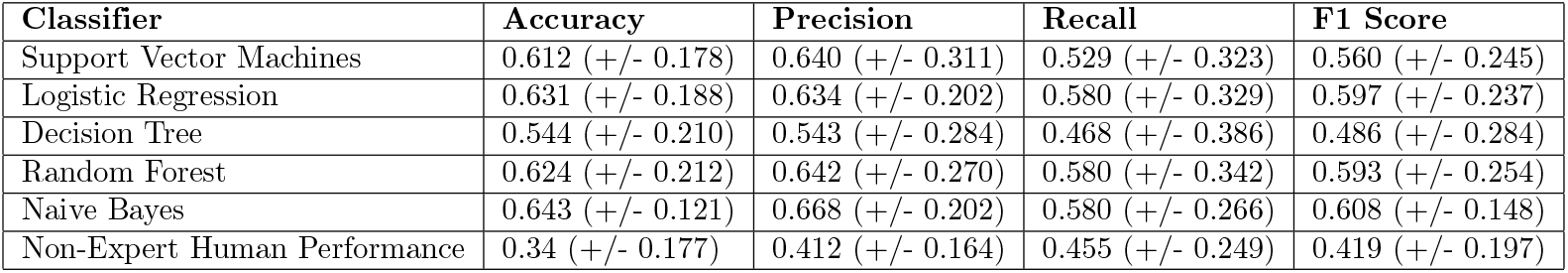
Evaluation of the classification performance of the tool with original video data from the wet-bench experiments. We used a cross-validation style test with 10 folds of classifications. This means that the classification experiment was run 10 times, varying the training and validation data. Each run used 90% of the data (144 videos) for training and 10% of the data (16 videos) for testing. Reported error is standard deviation.

| Classifier | Accuracy | Precision | Recall | F1 Score |
| --- | --- | --- | --- | --- |
| Support Vector Machines | 0.612 (+/- 0.178) | 0.640 (+/- 0.311) | 0.529 (+/- 0.323) | 0.560 (+/- 0.245) |
| Logistic Regression | 0.631 (+/- 0.188) | 0.634 (+/- 0.202) | 0.580 (+/- 0.329) | 0.597 (+/- 0.237) |
| Decision Tree | 0.544 (+/- 0.210) | 0.543 (+/- 0.284) | 0.468 (+/- 0.386) | 0.486 (+/- 0.284) |
| Random Forest | 0.624 (+/- 0.212) | 0.642 (+/- 0.270) | 0.580 (+/- 0.342) | 0.593 (+/- 0.254) |
| Naive Bayes | 0.643 (+/- 0.121) | 0.668 (+/- 0.202) | 0.580 (+/- 0.266) | 0.608 (+/- 0.148) |
| Non-Expert Human Performance | 0.34 (+/- 0.177) | 0.412 (+/- 0.164) | 0.455 (+/- 0.249) | 0.419 (+/- 0.197) |

Using the original trajectory images created from the videos as a statistical basis, we simulated an expansive trajectory feature space by fitting Gaussian distributions over the original features via a Gaussian Mixture Model (GMM). The goal was to generate as much evidence as possible for classifier inference. In this context, a GMM can be used as a generative probabilistic model that describes the distribution of the data and finds a mixture of multi-dimensional Gaussian probability distributions that best describe any input feature space. The appropriate parameters are subsequently fit to the GMM model to generate *n* synthetic feature vectors for each class. The value *n* = 10000 was selected via empirical observation. Results for different values of *n* can be seen at Supp. Tables 1 and 2. Table 2 shows the result after augmenting the training data via this simulation. As can be seen, the result of classification is drastically improved when more data is used. The best achieved result of 86.7% from the decision tree classifier is significantly higher than non-expert human performance, and brings the tool to a point of functionality that is useful to the experimentalist in practice.

**Table 2.** Evaluation of tool with simulated data (number of synthetic data points *n* = 10000) obtained after generative sampling using two different GMM models: one for positive samples, and one for negative samples. We used cross-validation for classification over 10 folds, meaning that for each fold 18000 samples are used for training and the remaining 2000 are used for testing. Reported error is standard deviation.

| Classifier | Accuracy | Precision | Recall | F1-score |
| --- | --- | --- | --- | --- |
| Support Vector Machines | 0.840 (+/- 0.019) | 0.842 (+/- 0.035) | 0.837 (+/- 0.019) | 0.839 (+/- 0.016) |
| Logistic Regression | 0.764 (+/- 0.016) | 0.772 (+/- 0.017) | 0.751 (+/- 0.021) | 0.761 (+/- 0.017) |
| Decision Tree | **0.867** (+/- 0.011) | **0.869** (+/- 0.008) | **0.863** (+/- 0.022) | **0.866** (+/- 0.013) |
| Random Forest | 0.707 (+/- 0.023) | 0.730 (+/- 0.030) | 0.659 (+/- 0.046) | 0.692 (+/- 0.029) |
| Naive Bayes | 0.712 (+/- 0.020) | 0.716 (+/- 0.022) | 0.702 (+/- 0.028) | 0.709 (+/- 0.021) |

## Discussion

Prior research suggests that cross-modal sensory integration occurs in all vertebrate species, including humans [7, 11, 50]. Any defects in this operation can cause serious neurological malfunctions, such as difficulties in positional awareness (proprioception), movement (vestibular system), or response to stimulation in various sensory systems [22, 27]. Tools that assist in the study this phenomenon, such as the one introduced in this paper, have the potential to accelerate clinical work in the future.

In almost all vertebrate species previously studied (e.g., teleost, reptiles, birds, rodents, primates), the brain sends centrifugal signals to the neural retina. Depending on the species, the centrifugal input may come from different regions of the brain. In monkeys, cortex projections to the lateral geniculate nucleus (LGN) have been discussed thoroughly by Briggs and Usrey [6]. That study revealed that the cortex signals sharpen/refine the receptive fields of LGN neurons and amplify signal transmission through the brain, thereby aiding directed attention. Further, the neural retinas receive histamine input from the hypothalamus [16]. The presence of histamine has been found to change membrane potentials of retinal bipolar cells and the firing patterns of retinal ganglion cells [1, 17]. In humans, prior work [18, 21] suggests sensory signaling interactions between the olfactory and visual systems. Imaging and behavioral analyses revealed the source of interactions to be in the forebrain, the exact location being somewhere between the anterior hippocampus and rostromedial orbitofrontal cortex. All of this research points to a common theory that suggests that measures of performance or behavior in response to sensory stimulation are amplified, particularly when a stimulus in one modality is ambiguous or under-determined. The molecular and cellular pathways involved in such types of sensory integration, however, remain an active area of investigation.

The zebrafish is a standard model system for vision studies because its retina is strikingly similar to that of mammals [28, 47]. Sensory integration in zebrafish is mediated by the olfacto-retinal centrifugal pathway. The application of odors (e.g., by administration of the amino acid methionine) activates the olfactory sensory neurons in the olfactory epithelia, thereby increasing the release of the neurotransmitter glutamate in the olfactory bulb. The TN, located in the olfactory bulb, expresses glutamate receptors, and is thus activated after odor stimulation. The TN synthesizes and releases the hormone GnRH, and synapses with dopaminergic cells in the retina. The neurotransmitter dopamine inhibits retinal ganglion cell activity. Activation of GnRH inhibits dopamine release from dopaminergic cells in the retina. At the end, due to olfactory stimulation, dopamine content in the retina is decreased, and the inhibition of dopamine to retinal ganglion cells is lifted. Thus visual sensitivity will be increased. A key to increasing understanding of this cross-modal sensory integration is the ability to conduct experiments at scale, with computer-assisted analysis.

When it comes to building computational models for behavior analysis, Niu et al. [38] provide a comprehensive survey of recent studies analyzing fish behavior through computer vision. The fish behaviors described in this study include schooling, swimming, stress response, feeding, taxis, reproduction and migration. Different behaviors have different performance characteristics and implications. For example, monitoring fish swimming behavior and stress helps scientists to uncover problems related to overall climatic and environmental changes that affect all of us.

Most similar to our research are works that study how to detect and track multiple zebrafish with frequent occlusion for behavior analysis [2, 41, 42, 49]. Bai et al. [2] suggest the use of the histogram of oriented gradients (HOG) as a feature representation for identifying individual fish, while Xu et al. [49] use a shallow convolutional neural network. Qian et al. [41] use fish head detection via extremum detection and ellipse fitting, along with Kalman filtering with feature matching, to detect and track individual fish in complex motion. Their other work [42] uses a complex combination of fish head detection and tracking via moving region segmentation, centerline extraction, head direction estimation and global optimization association. Our work is different from the existing research in the sense that our proposed tool does not end with tracking, but shows how we can utilize the information we gain from the tracker for automatic classification of behavioral changes. We develop a fully automatic end-to-end pipeline utilizing state-of-the-art deep learning methods for detection and a straightforward method for the tracking of fish. Occlusion does not affect our method as much as the others since we utilize it to our advantage by treating it as a form of noise for training to help the autoencoder, which produces the features for classification, generalize. In comparison to other methods described above, our method does not rely on elaborate calculation of head poses and only uses a fraction of the amount of labelled data for training the detection and classification modules due to data augmentation via the GMM. Hence, at test time, it is quite fast and accurate. Analyzing complex behavior phenotypes, as reflected in responses to stimuli (external or internal) triggered by signal transduction in sensory neurons, interneurons, and motor neurons in the spinal cord as well as in the brain cortex, would be hard for our tool to differentiate. However, if such videos and their corresponding labels are available beforehand, our classifier module can be fine-tuned to reflect those fine-grained classes. The task of differentiating behavioral changes in fish is difficult (as shown by our user study) and hence, cannot be easily crowdsourced. Thus our tool can be used by biologists as-is or as a pre-processing step to skim through videos to surface the most interesting ones for more exhaustive manual analysis. By re-training the learning-based components (e.g., the detection module to identify animals other than fish), our tool can also be used to study other motion-based animal behavior.

## Methods

### Procedure for Behavioral Experiments

Previously, we developed a behavioral assay based on visually-mediated escape responses to measure the behavioral visual sensitivity in zebrafish (*Danio rerio*) [29]. The animals used in this research were between 4-6 months old, both male and female. The fish were maintained in 28C circulating water (ph 7.0) under 14/10 light/dark cycles (ceiling light). They were fed two times a day with freshly hatched brine shrimp. The test apparatus (see Fig. 2(A)) consisted of a transparent container surrounded by a rotating drum. The drum was illuminated by a light source from above, and the light intensity could be adjusted by changing neutral density filters. The drum could rotate in either clockwise or counterclockwise directions. The inside of the drum was covered by white paper marked with a black segment. The fish was allowed to freely swim in the container, and a post was placed in the middle of the container to prevent the fish from swimming through the center of it. The swimming behavior was viewed via a monitor connected to an infrared video camera. Normally, the fish swam slowly along the wall of the container in either a clockwise or counterclockwise direction. However, when challenged by the black segment rotating outside the container, the fish displayed robust escape responses, i.e., as soon as the black segment came into view, the fish immediately turned and rapidly swam away. By measuring the minimum light intensities required to evoke the escape responses, we evaluated the visual sensitivity of zebrafish. The experiments were conducted as follows:

1. The fish was transferred to the test container, one fish per container.
2. The fish was dark adapted in complete darkness for 30 minutes.
3. The fish was tested for behavioral responses to the approach of the rotating segment. Initially, the intensity of light that illuminated the rotating segment was set at a near complete dark level (log I = −6.0; the maximum light intensity measured at log 0 = 425*μW/cm*^2^), then gradually increased (by removing neutral density filters) at 0.5 log unit steps until the fish showed escape responses to the rotating segment. The minimum light required for eliciting escape responses was noted as the threshold light sensitivity (the absolute visual sensitivity level).
4. Upon determining the light threshold, odor stimulation (1 microliter stock solution of methionine, 3mM dissolved in water) was administrated to the test container via a pipette.
5. Immediately after the administration of methionine, the visual sensitivity of the fish was measured again.
6. Repeat steps 1 – 3.

### A Tool to Analyze Behavior

Our tool builds on the flexibility and success of deep learning methods for detection and optical flow for tracking. The entire pipeline consists of five main stages: (1) automatic fish detection, (2) tracking of fish, (3) compression of trajectory images for further data generation, (4) data augmentation through generative sampling and (5) binary classification to determine if a behavioral change is present. The process begins with a raw video showing the movement of fish in water after the application of chemicals and ends with a binary “yes” or “no” prediction indicating whether the fish exhibited behavioral change or not.

Since the behavioral change in fish is primarily learned through its swimming pattern, we use automatic target detection and tracking in parallel to generate a trajectory image from a single video. Effectively, the trajectory images holds information about the swimming pattern of a fish before and after the application of odorants in water and is created by fusing images acquired as a result of detection and tracking from videos. Since these trajectory images can be large, we use compression via an autoencoder model to reduce the number of features before sending them to a classifier. We then use the features acquired after compression of the trajectory images to classify whether or not a fish shows any behavioral change. Most machine learning methods are data hungry and cannot generate trustworthy results if sufficient amounts of data are not available. With only a relatively small number of videos is available from the wet-bench experiments, we simulate an expansive data space by sampling from a GMM over the original data. The goal is to generate as much evidence as possible for statistical inference. In the following subsections, we provide the details about each stage.

#### 1. Detection

A target detection algorithm is employed to locate the fish in each frame. The detector is primarily employed to create a mask for fish and apply it on individual frames before the optical flow operation, such that only the displacement of fish between frames is captured by the optical flow algorithm, ignoring the motion of other pixels in the frame representing the apparatus for the experiment. For automatic detection, we use YOLOv3 [43], a deep convolutional neural network for real-time object detection at various scales. Since YOLOv3 is primarily trained on the PASCAL VOC [13] and MS-COCO [32] datasets, we re-trained it specifically on our fish dataset to identify fish under very little light. This dataset is made available to researchers for further work on this problem (see Data Availability Statement). Compared to its previous version YOLOv2, YOLOv3 has a deeper architecture and contains residual connections and upsampling layers. Some salient features of YOLOv3 that are improvements over the previous versions are its ability to make detections at three different scales and to detect smaller objects, both of which make it useful for detecting fish in tanks. The algorithm outputs bounding box coordinates for a detected fish. The centroid of the bounding box is then calculated to locate the middle of the fish, with a circle with a 10 pixel radius as the point representing the location of fish in the tank. This point serves as a key point for the optical flow algorithm. The motivation for using a point instead of a rectangular bounding box came from the observation that it is easier to create a trajectory image via points.

#### 2. Tracking

For tracking we use a dense optical flow method [15] that shows the displacement of each and every pixel between frames. The algorithm operates on two corresponding frames as inputs to produce a single image that shows the displacement. We use OpenCV’s implementation of Farnebäck dense optical flow [15] for tracking the fish across video frames. The algorithm begins by approximating the windows of video frames by quadratic polynomials through polynomial expansion transform. Then, by observing how the polynomial transforms under motion, a method to estimate displacement fields from polynomial expansion coefficients is defined. After a series of refinements, dense optical flow is finally computed. For example, for a particular video of *x* seconds with *n* + 1 number of video frames, we use the detection algorithm for *n* + 1 frames and end up with masks for each frame, representing the location of the fish. We then use these masks for tracking via optical flow, which results in *n* frames showing the displacement of fish between frames. Next, we combine these *n* frames to generate a single trajectory image for the entire video. The process of creating the trajectory image is illustrated in Fig. 2(B). Note that this optical flow method is highly dependent on the success of the detection algorithm that is applied at the previous stage.

#### 3. Trajectory Image Compression

Since the raw features of a trajectory image are large (256 × 256 × 3) in comparison to the number of videos collected from the wet-bench experiments (160), we need to compress them before generative sampling and classification. We use an autoencoder for that purpose. The autoencoder [37] is a standard approach for learning compact object representations and is widely used as a data compression algorithm where the compression and decompression functions are data-specific, lossy, and learned automatically from examples. It takes an input *x* ∈ [0, 1]^*d*^ and first maps it with an encoder to a hidden representation *y* ∈ [0, 1]^*d*′^ through a deterministic mapping: *y* = *s*(*Wx* + *b*) where *s* can be a non-linearity like the sigmoid or RELU functions. The latent representation is then mapped back into a reconstruction *z* of the same shape as *x*, i.e., *z* = *s*(*W′y* + *b′*). The reconstruction error can be measured through traditional squared error. If the input is interpreted as either bit vectors or vectors of bit probabilities, cross-entropy of the reconstruction can be used. For the purpose of our task, we used the convolutional layers of VGG19 [45] and compressed the raw features to a size of 64 dimensions. These features can then be directly used for classification (see Fig. 2(C) for a diagram of the entire process) or can be used to generate synthetic data for improving classification.

#### 4. Data Augmentation Via Generative Sampling

Most supervised machine learning algorithms depend on large amounts of data for training. Since we are operating on a relatively limited number of videos from wet-bench experiments, we simulated an expansive dataset for training the classifier stage of the tool using a GMM. A GMM is fundamentally a probabilistic model that assumes all of the data points are generated from a mixture of a finite number of Gaussian distributions with unknown parameters. The approach is often used for classification and generative modeling, as well as for multimodal data. This is because it provides a richer class of density models than a single Gaussian distribution. A limitation of the GMM approach is that the loss function is non-convex and optimizing it is non-trivial. The most popular algorithm that is used for optimizing GMMs is the Expectation Maximization algorithm [34].

The GMM is parameterized by two types of values: the mixture component weights and the component means and variances/covariances. For univariate data analysis, a GMM is defined as a linear superposition of *K* components by:

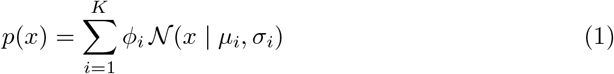

where *μ_i_* and *σ_i_* represent the mean and variance of the *i*th component, *ϕ_i_* represents the mixture component weight for component *C_i_*, with the constraint that 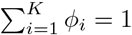 so that the total probability distribution normalizes to 1. 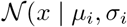 represents the individual normal distribution for component *C_i_*. Since the GMM is completely determined by the parameters of its individual components, a trained GMM can give an estimate of the probabilities of both in-sample and out-of-sample data. Moreover, since we can numerically sample from individual Gaussian distributions, we can sample from a GMM in a straightforward manner to create synthetic datasets.

For our task, we use a GMM to model the overall distribution of the input data. We assumed that the entire trajectory space can be represented as a mixture of Gaussian distributions. Some of the motivating factors that led us to use GMMs in lieu of other generative sampling methods such as GANs [20] were limited samples, dimensionality of the trajectory images, and the simplicity of the calculation. In comparison to other unsupervised density estimation methods, GMMs can operate with limited data. Since there can be two outcomes of our task, i.e., the fish exhibits a behavioral change after the application of the odorant into the water, or there is no change in behavior, we train two separate GMM models from our training data, i.e., the features obtained after compressing the original 160 videos from from the wet-bench experiments with the autoencoder. From these two GMM models, 10000 feature vectors are generated from each one for classifier training, which is discussed in the next section.

#### 5. Binary Classification for Predicting Behavioral Changes

Here we ask the following question: given a set of features representing a trajectory image after compression, is it possible to identify whether any changes in the movement of a fish, as encoded by the features, has been triggered after an olfactory signal? Mathematically, if *D* =(*x_i_*, *y_i_*) of size *n* represents the dataset of sampled generated trajectory features after compression, *x_i_* being the trajectory features such that *x* ∈ *R^n^* and *y_i_* ∈ [0, 1] being the corresponding labels for behavior change (1) or otherwise (0), the task of identifying the fish behavior from a new compressed trajectory feature, *x_new_*, can be expressed as a function *f_θ_*(*x_new_*) parameterized by *θ* after being trained on *D*, given by the following expression:

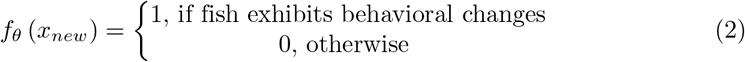

In essence, this task can be formulated as a binary classification problem. Ideally, any discrimitative supervised learning method can be employed to solve the problem. We use Support Vector Machines, Decision Trees, Logistic Regression, Naive Bayes and Random Forest as possible classification candidates, and find that Decision Trees yield the best outcome (see Results Section).

##### Support Vector Machines

The Support Vector Machine (SVM) is a supervised learning paradigm that is widely used in classification and regression tasks [9]. Since the features obtained as a result of the compression of the trajectory images are numeric and high-dimensional, we use a linear SVM formulation, which is suitable for such data.

An SVM classifier utilizes a subset of training points, commonly referred to as “support vectors”, in the decision function to define the decision boundary between classes. During training, it attempts to find the optimal hyperplane for classifying test samples based on these support vectors and some constraints. Given a training dataset, *D* = (*x_i_*, *y_i_*) of size *p* with *x_i_* = (*x*_*i*,1_, *x*_*i*,2_,…, *x_i,q_*) and label *y_i_* = −1 or +1, formally the SVM classifier can be defined as a quadratic optimization problem solving the following equation:

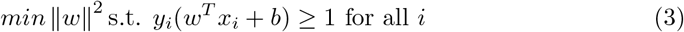

where *w* = (*w*_1_, *w*_2_,…, *w_q_*) represents the weight vector and *b* the bias. An important point to be noted when training an SVM model is the parameter *C* that controls the trade-off between having a wide margin and correctly classifying training data.

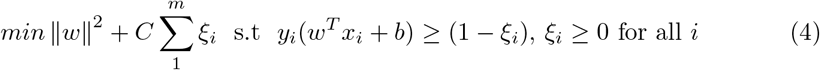

A larger value of *C* indicates a smaller number of mis-classified training samples and is susceptible to overfitting.

##### Gaussian Naive Bayes

Another common method that is often used for supervised classification tasks is Naive Bayes [48]. Naive Bayes is a very simple probabilistic classification algorithm that makes strong assumptions about the independence of each attribute of the data and is based on the classical Bayes’ theorem. Naive Bayes calculates the posterior probability of an observation *v* belonging to a class *C_k_*, *p*(*C_k_*|*v*) by combining the likelihood of observation *v* belonging to the class *C_k_*, *p*(*v*|*C_k_*) and prior probabilities of *C_k_*, *p*(*C_k_*).

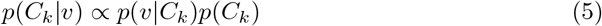

Gaussian Naive Bayes extends the concepts of simple naive Bayes to that of real-valued continuous attributes. For example, if *x* represents a real-valued attribute, *μ_k_* is the mean and 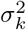 the variance of *x* associated with the the class, *C_k_*. Then, for Gaussian Naive Bayes, the likelihood or probability of some observation *v* given the class *C_k_*, *p*(*x* = *v*|*C_k_*) can be computed by inserting *v* into the equation for a Gaussian distribution parameterized by *μ_k_* and 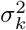:

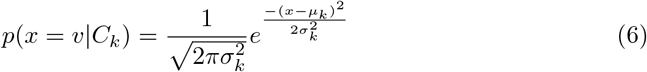

Now if *x* = (*x*_1_,*x*_2_,..,*x_n_*) is a vector representing *n* independent features, the likelihood would be given by 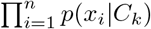 and the Naive Bayes classifier for class *ŷ* = *C_k_* for some *k* is defined as:

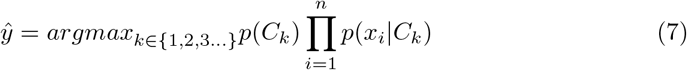

In the present context, *x* stands for the features acquired after compressing the trajectory image and the classes *C_k_, k* = 0, 1 are binary (0 for no behavioral change and 1 for noticeable changes in behavior).

##### Logistic Regression

Logistic regression belongs to the family of generalized linear models that are commonly used to model a binary categorical variable using numerical and categorical predictors. It assigns weights to each input attribute or feature and outputs a value between 0 and 1. This output can be viewed as a probability of success relative to the target variable, with any probability ≥ *p, p* ≥ 0.5 being considered a “success”.

Given a set of instance-label pairs (*x_i_, y_i_*), where *x* ∈ *R^n^, i* = 1, 2,…*l* and *y* ∈ {+1, −1}, logistic regression (L2- regularised) solves the following unconstrained optimization problem with loss function, *ξ*(*w*; *x_i_, y_i_*) [14]:

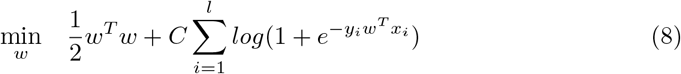

where *C* > 0 is a penalty parameter, and *w* represents the weight vector. Similar to Naive Bayes, for our task, *x_i_* stands for the features acquired after compressing the trajectory image and the class *y_i_* is binary (0 for no behavioral change and 1 for a noticeable change in behavior).

##### Decision Tree

Decision trees are one of the most popular supervised and computationally inexpensive machine learning tools for classification when the data is continuously split into two or more homogeneous subsets according to a certain parameter. It primarily consists of the following structures: a `root` node that has no incoming edges and zero to multiple outgoing edges. `internal` nodes each of which has exactly one incoming edge and multiple outgoing edges. `edges or branches` that connect between nodes. `leaf or terminal` nodes each of which has exactly one incoming edge and no outgoing edges. Typically, the leaves represent class labels and branches represent conjunctions of features that lead to those class labels. The non-terminal nodes (root and internal nodes) contain attribute test conditions based on which data are split or separated due to different characteristics. The criteria for separation can be calculated through information gain or entropy calculation. It works for both categorical and continuous input and output variables. Decision trees are notorious for over-fitting their data. Methods to overcome this include carefully pruning the tree, using early stopping, or ensemble methods.

##### Random Forest

A random forest [5] is a supervised ensemble method consisting of a number of decision trees. It aggregates the votes from different decision trees to decide the final class of the test object in order to improve the predictive accuracy and control over-fitting. It is common to find that individual decision trees within the random forest exhibit high variance and tend to overfit. To decrease variance and control over-fitting, each tree is built from a sample drawn with replacement from the training set. Moreover, when splitting each node during the construction of a tree, the best split is found either from all input features or a random subset of maximum features. This process of injecting randomness provides individual decision trees with decoupled prediction errors, and by taking an average of the predictions, some errors cancel out, yielding a better model.

### Crowdsourced Data Annotation Study

To compare the effectiveness of our newly introduced tool against human performance and to check whether the task can be crowdsourced to the untrained public or not, we conducted a user study (see Supp. Fig. 3). The complete procedure for the study is as follows. A participant is first presented with a video clip *a* positioned on the top of a screen, labelled as the “Before” (pre-treatment) video and another video clip *b* right below the first, labelled as the “After” (post-treatment) video. The observer is informed that *a* and *b* represent the fish swimming in the experimental apparatus before and after the application of chemicals (odorant) in water respectively. Below the video pair, three options are provided and the observer is asked to select the option that most applies to the question, “Do you think the fish reacts differently after being treated with chemicals?”

To capture as much of the underlying complexities in human judgment as possible and to aid in decision making, a list of common behavioral traits is provided to the subjects. Examples of those traits include turn and follow response, i.e., the fish changes swimming pattern in order to follow the stimulus, showing signs of interest in the stimulus; escape response (the fish turns and swims in the opposite direction after becoming aware of the presence of the stimulus); dodging response (the fish jumps or swims away from the stimulus, towards the middle of the tank); flinches or jumps (the fish jumps or flinches slightly away from the stimulus but doesn’t change swimming pattern); changes in speed (the fish speeds up or slows down upon acknowledgement of the stimulus, but does not change direction of swimming). Each subject is given unlimited time and is informed that providing an accurate assessment is most important.

A total of 10 pairs of pre- and post-treatment videos were selected for the study. 11 subjects participated in the study, consisting of college students and professionals who were completely unfamiliar with the process of behavioral studies in neuroscience. Because of the lack of familiarity the subjects had for the task, we included a small tutorial detailing the process within the study. The intention of using such novice participants stemmed from the possibility of crowdsourcing these types of data analysis tasks in neuroscience on platforms like Amazon’s Mechanical Turk service, where participants at the level of competency of our test subjects are abundant and the cost of annotation is very low.

## Acknowledgements

The research was funded by the Department of Defense (Army Research Laboratory) under the contract W911NF-18-1-0292.

## Author contributions statement

S.B. designed, analyzed, and implemented the model, and led the writing of the paper. L.L. designed the wet-bench experiments; L.A. and P.B. were responsible for conducting the wet-bench experiments and preparing the source data. S.Y. helped in the literature survey. W.J.S. supervised the tool development and helped in writing the manuscript.

## Data Availability Statement

The datasets analyzed for this study and the source code used for modeling have been released for reproducability and can be downloaded via https://github.com/sbanerj2/zebrafish_behavior.

## Additional information

All experiments involving animals were carried out in accordance with the guidelines of the National Institutes of Health, and approved by University of Notre Dame IACUC (Protocol #17-11-4243). The crowdsourced data annotation study (see the Methods section) involving human participants has been approved by the University of Notre Dame IRB under protocol #18-01-4341. Informed consent was obtained from all participants involved in this study. The study was carried out in accordance with relevant guidelines.

