## Supplementary material for "An Assistive Computer Vision Tool to Automatically Detect Changes in Fish Behavior In Response to Ambient Odor": Supp. Table 1, Supp. Table 2, Supp. Fig 1, Supp. Fig 2, Supp. Fig 3 (from another version of this preprint: version 1, 2020-09-02)

#### Detection Results

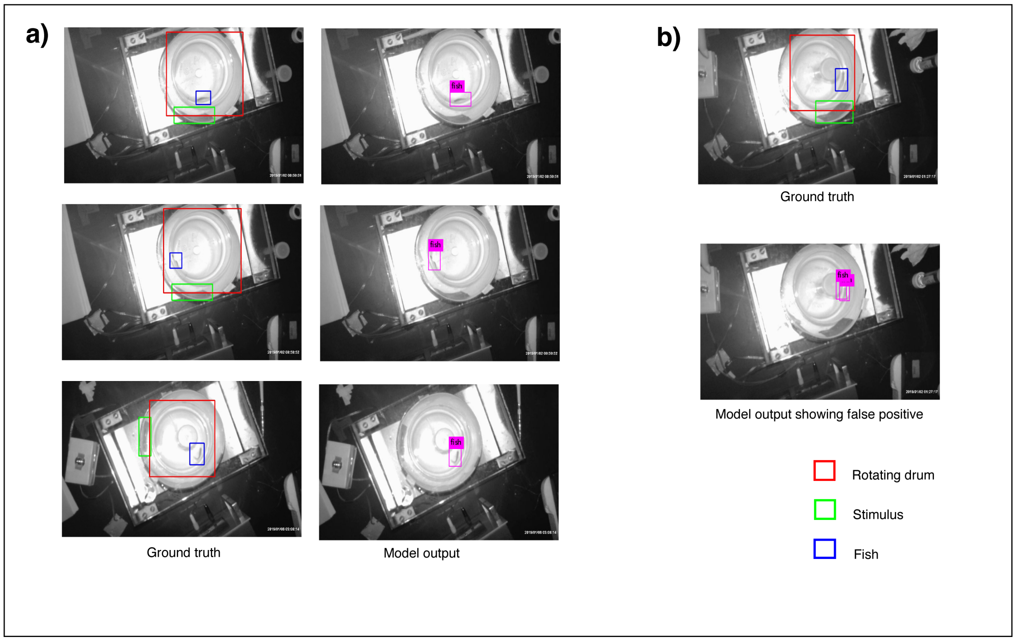


**Figure 1. Detection Results.** (a) Positive detection results where the fish is correctly identified within the tank. (b) A failure case for the detector where it incorrectly identifies the reflection of the fish within the tank as another fish. The ground-truth annotation is given on the left for (a) and top for (b). The blue, red and green rectangles (or bounding boxes) in the ground-truth images represent the fish, the moving cylinder and stimulus. Most fish move away or towards the stimulus.

#### Example reconstructed outputs from different autoencoder models

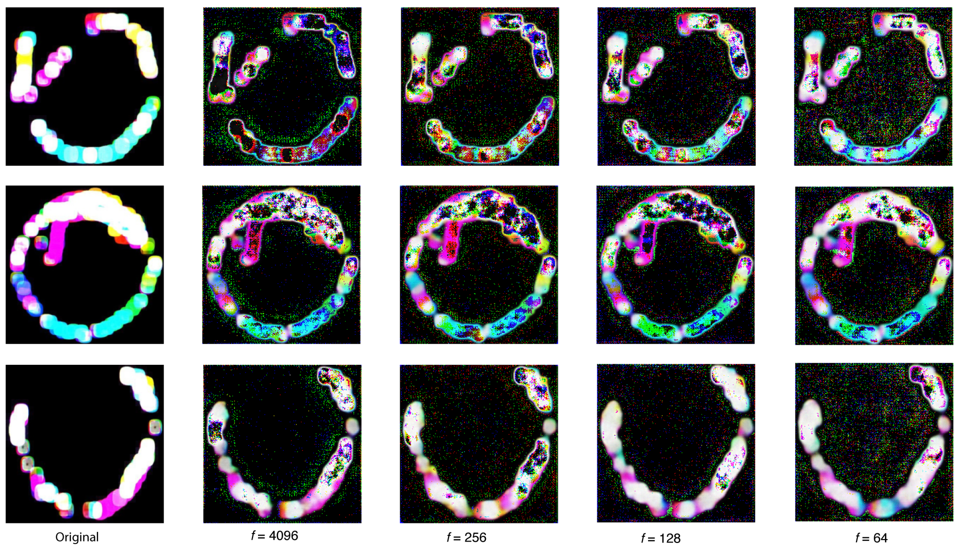


**Figure 2. Example reconstructed outputs from different autoencoder models:** Here $f$ represents the dimensionality of the latent representation from the autoencoder, i.e., the encoded feature dimensionality. A larger feature dimensionality usually results in a better reconstruction.

### Choice of number of GMM samples

| Classifier | Accuracy | Precision | Recall | F1-score |
|---|---|---|---|---|
| Support Vector Machines | **0.822** (+/- 0.036) | **0.835** (+/- 0.052) | **0.802** (+/- 0.050) | **0.818** (+/- 0.036) |
| Logistic Regression | 0.749 (+/- 0.062) | 0.763 (+/- 0.086) | 0.725 (+/- 0.055) | 0.743 (+/- 0.057) |
| Decision Tree | 0.770 (+/- 0.065) | 0.775 (+/- 0.065) | 0.760 (+/- 0.077) | 0.767 (+/- 0.067) |
| Random Forest | 0.680 (+/- 0.059) | 0.684 (+/- 0.072) | 0.672 (+/- 0.102) | 0.677 (+/- 0.063) |
| Naive Bayes | 0.702 (+/- 0.059) | 0.706 (+/- 0.062) | 0.697 (+/- 0.116) | 0.700 (+/- 0.070) |

**Table 1.** Evaluation of tool with simulated data (number of synthetic data points $n = 2000$) obtained after generative sampling using two different GMM models: one for positive samples, and one for negative samples. We used cross-validation for classification over 10 folds, meaning that for each fold 1800 samples are used for training and the remaining 200 are used for testing. Reported error is standard deviation. The results in this Table are lower than those reported in Table 2 in the main article.

| Classifier | Accuracy | Precision | Recall | F1-score |
|---|---|---|---|---|
| Support Vector Machines | **0.715** (+/- 0.195) | **0.746** (+/- 0.211) | 0.660 (+/- 0.360) | **0.689** (+/- 0.239) |
| Logistic Regression | 0.620 (+/- 0.211) | 0.638 (+/- 0.264) | 0.580 (+/- 0.307) | 0.600 (+/- 0.232) |
| Decision Tree | 0.560 (+/- 0.117) | 0.583 (+/- 0.193) | 0.510 (+/- 0.140) | 0.537 (+/- 0.092) |
| Random Forest | 0.580 (+/- 0.206) | 0.600 (+/- 0.264) | 0.490 (+/- 0.316) | 0.529 (+/- 0.273) |
| Naive Bayes | 0.640 (+/- 0.199) | 0.636 (+/- 0.193) | **0.680** (+/- 0.294) | 0.650 (+/- 0.210) |

**Table 2.** Evaluation of tool with simulated data (number of synthetic data points $n = 200$) obtained after generative sampling using two different GMM models: one for positive samples, and one for negative samples. We used cross-validation for classification over 10 folds, meaning that for each fold 180 samples are used for training and the remaining 20 are used for testing. Reported error is standard deviation. The results in this Table are much lower than those reported in Table 2 in the main article.

### Human Study

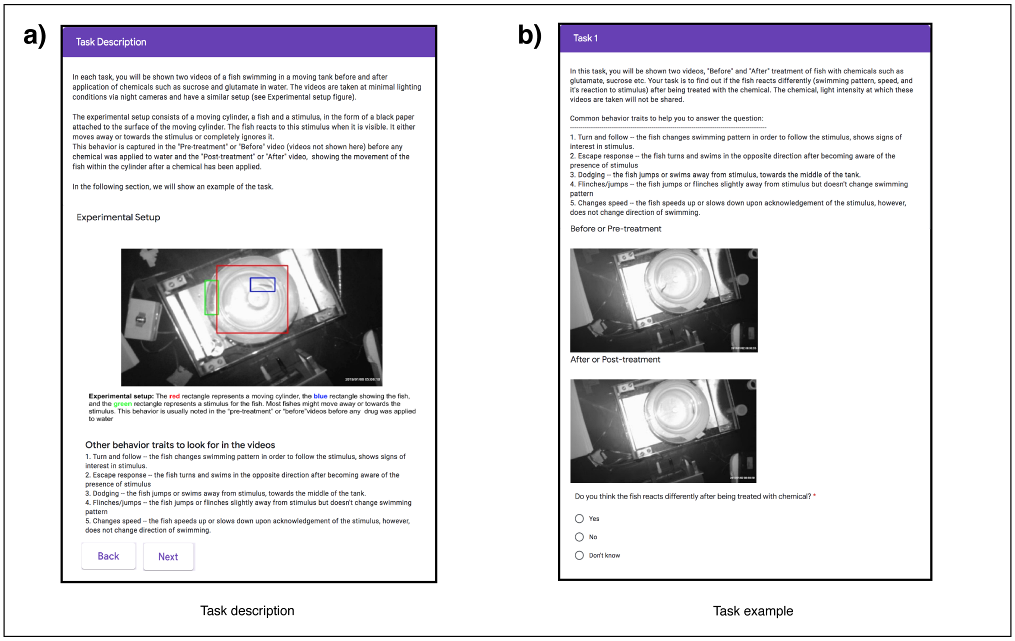


**Figure 3. Screenshot of the human study meant to evaluate the effectiveness of crowdsourcing the video analysis task to non-experts.** (a) A task description along with an example was provided to non-experts in order to aid them in the analysis of videos showing fish behavioral changes. (b) An example of the actual task the participants were given. Note that the actual pre- and post-treatment videos have been replaced by still images in this figure.
